## Supplementary material for "Essentiality of c-di-AMP in *Bacillus subtilis*: Bypassing mutations converge in potassium and glutamate homeostasis": S1

**S1 Supporting Information**

**Global gene expression in the *B. subtilis* wild type strain in response to the potassium concentration and the nitrogen source.**

In order to study the consequences of changing environmental potassium and glutamate concentrations, we cultivated *B. subtilis* 168 in MSSM minimal medium with 0.1 or 5 mM KCl and with ammonium or glutamate as the nitrogen source, and analyzed the global transcriptomes.

First, we considered the expression of the genes involved in c-di-AMP homeostasis (S2 Table). Of the three genes encoding the diadenylate cyclases, *cdaA* and *disA* were expressed under all tested conditions, whereas the *cdaS* gene was not expressed with ammonium as the nitrogen source or with glutamate at a low potassium concentration. Even in the presence of 5 mM potassium and glutamate, the expression of *cdaS* was very low as compared to the other cyclase-encoding genes. The expression of *cdaA* and *disA* was about three-fold reduced at the high potassium concentration if glutamate was present. In contrast, these genes were not affected by the potassium concentration when ammonium was the nitrogen source. The *gdpP* and *pgpH* genes that encode the two c-di-AMP-degrading phosphodiesterases were also expressed under all tested conditions. Again, the expression was reduced at 5 mM potassium if glutamate was present as the nitrogen source. The fact that the diadenylate cyclase and phosphodiesterase genes are regulated in the same manner, even though the encoded enzymes catalyze opposing reactions, suggests that the control is mainly exerted at the post-transcriptional level.

In the presence of glutamate, the *kimA* and *ktrAB* genes encoding high-affinity potassium transporters were 112- and 29-fold induced by potassium limitation, whereas the *ktrC* and *ktrD* genes encoding the low-affinity potassium transporter were not affected by the potassium concentration. With ammonium as the nitrogen source, the *kimA* and *ktrAB* genes were much less responsive to potassium availability (17- and 4-fold induction by potassium limitation, respectively). Interestingly, the expression of all potassium transporters is increased in the presence of glutamate if potassium is limiting in the medium. The expression of the *gltAB* operon encoding the glutamate synthase is 8- and 33-fold repressed by glutamate in the presence of 0.1 mM and 5 mM KCl, respectively. Importantly, in the absence of glutamate, the expression of the operon is fivefold enhanced at the increased potassium concentration. Thus, the gene expression data confirm the interrelation between potassium and glutamate homeostasis. To obtain independent evidence for this hypothesis, we studied the activities of the *kimA* and *gltAB* promoters using fusions of a promoterless *lacZ* gene encoding β-galactosidase to the corresponding promoter regions. For the *kimA-lacZ* fusion present in *B. subtilis* GP2181, we observed a strong repression at increasing KCl concentrations, as described earlier (Gundlach *et al*., 2017). In addition, the expression of *kimA* was strictly dependent on the presence of glutamate, even at low potassium concentrations (see S2 Fig.). The expression of the *gltA-lacZ* fusion was assayed using strain GP342. In the absence of glutamate, glutamate synthesis, and thus expression if the *gltAB* operon, is essential for the bacteria. However, the activity of the *gltA* promoter is increased from 300 to about 450 units per mg of protein, if the potassium concentration in the medium is increased from 0.1 mM to 5 or 20 mM KCl (S2 Fig.). Thus, the reporter assays support the conclusion that the expression of potassium transporters is increased in the presence of glutamate whereas the expression of the *gltAB* operon is enhanced at high potassium concentrations. These results are in excellent agreement with the observed interdependence of the cellular potassium and glutamate pools, and provide an explanation for this mutual dependence of the two ions.

We also analysed the transcriptome data to identify the genes and operons that are most strongly affected by the availability of potassium and glutamate. In addition to the potassium transporters, genes of the Rex regulon that are involved in respiration and fermentation are strongly repressed by potassium (in particular in the absence of glutamate): the *cydABCD* and the *ldh-lctP* operons encoding the high-affinity terminal quinol oxidase and the lactate dehydrogenase are repressed 563 and 327-fold, at increased potassium concentrations (see S1 Table). In contrast, most sporulation genes are switched off at limiting potassium concentrations suggesting an increased demand for potassium in spores. Finally, the genes of the Fur regulon that are normally induced upon iron starvation depend on potassium for their expression. The availability of glutamate, as expected, most drastically affected the genes of the TnrA regulon that are required for the acquisition of alternative nitrogen sources. These genes are strongly repressed by the preferred nitrogen source ammonium. Interestingly, the *tapA-sipW-tasA* and *epsA-O* operons encoding the major components of the extracellular biofilm matrix are both strongly dependent on the presence of glutamate, and the expression is further enhanced if sufficient potassium (5 mM KCl) is available. The expression of these biofilm genes is not known to be regulated by any of the known regulators involved in nitrogen or potassium regulation. Finally, the Rex and SigD regulons involved in respiration and motility, are strongly repressed by glutamate in the presence of 0.1 mM or 5 mM KCl, respectively.

Gundlach J, Herzberg C, Kaever V, Gunka K, Hoffmann T, Weiß M, et al. Control of potassium homeostasis is an essential function of the second messenger cyclic di-AMP in *Bacillus subtilis*. Sci Signal. 2017; 10: eaal3011.

**S1 Table.**
**Transcriptomic data for the genes that are most strongly affected by the absence of c-di-AMP (Ammonium 0.1 mM KCl).** [1]

| Gene Name | Wild type | | | | GP2222 | GP2223 |
|---|---|---|---|---|---|---|
| | Glutamate 0.1 mM KCl | Glutamate 5 mM KCl | Ammonium 0.1 mM KCl | Ammonium 5 mM KCl | Ammonium 0.1mM KCl | Ammonium 5 mM KCl |
| Upregulated in Δ*dac* | | | | | | |
| *abrB* | 2656 | 2945 | 1141 | 2853 | 10065 | 15703 |
| *ndhF-ybcC* | 52 | 29 | 106 | 46 | 716 | 56 |
| *nrgAB* | 94214 | 140635 | 335 | 406 | 2064 | 793 |
| *glnRA* | 23015 | 10532 | 2487 | 8233 | 14422 | 18522 |
| Downregulated in Δ*dac* | | | | | | |
| *yoaH* | 1149 | 124 | 944 | 2844 | 206 | 957 |
| *lytF* | 3981 | 838 | 4508 | 23400 | 975 | 9865 |
| *yfmG* | 1575 | 9329 | 1116 | 2588 | 232 | 1657 |
| *hag* | 79726 | 10341 | 89536 | 289197 | 18386 | 183228 |
| *ybdO* | 1735 | 672 | 1539 | 7172 | 315 | 3364 |
| *epr* | 6635 | 557 | 5952 | 25153 | 1216 | 11613 |
| *bmrU-bmr-bmrR* | 319 | 325 | 1177 | 166 | 237 | 197 |
| *yqbG* | 160 | 28 | 132 | 342 | 26 | 94 |
| *pgdS* | 10133 | 1310 | 7581 | 26097 | 1453 | 7445 |
| *yolAB* | 8079 | 2632 | 6905 | 32510 | 1307 | 13367 |
| *yorG* | 32 | 26 | 124 | 18 | 23 | 16 |
| *ydaC* | 50 | 36 | 420 | 19 | 79 | 20 |
| *ykuNOP* | 28 | 452 | 150 | 1076 | 28 | 608 |
| *tlpC* | 650 | 250 | 386 | 1920 | 69 | 572 |
| *hemAT* | 3071 | 1448 | 2749 | 13874 | 477 | 3405 |
| *yfmTS* | 6151 | 794 | 6869 | 22920 | 1135 | 8665 |
| *sspF* | 150 | 1951 | 6456 | 351 | 996 | 455 |
| *yonX* | 34 | 153 | 300 | 17 | 46 | 18 |
| *mcpC* | 2789 | 605 | 5106 | 16420 | 770 | 6978 |
| *yscB* | 863 | 139 | 1373 | 4568 | 181 | 2124 |
| *ywfA* | 3525 | 3510 | 911 | 45 | 120 | 54 |
| *yxbCD* | 5251 | 21069 | 6976 | 6617 | 905 | 10137 |
| *araE* | 41 | 50 | 660 | 19 | 76 | 35 |
| *yxbBA-yxnB-asnH-yxaM* | 4359 | 36716 | 8577 | 10124 | 962 | 14183 |
| *yxkH* | 429 | 68 | 286 | 2268 | 30 | 457 |
| *sivC* | 5987 | 2337 | 9378 | 40055 | 899 | 25992 |
| *yorE* | 32 | 28 | 237 | 21 | 21 | 18 |
| *yvrJ* | 26 | 26 | 189 | 18 | 15 | 18 |
| *ywcJ* | 60 | 75 | 2449 | 42 | 163 | 75 |

| | | | | | | |
|---|---|---|---|---|---|---|
| *comGA---GG* | 19276 | 7644 | 545 | 294 | 31 | 63 |
| *yomK* | 758 | 71 | 880 | 1993 | 50 | 1324 |
| *sboAX* | 178 | 40 | 659 | 29 | 34 | 74 |
| *sigO-rsoA* | 65 | 46 | 661 | 70 | 28 | 97 |
| *tlpA-mcpA* | 668 | 141 | 911 | 1777 | 38 | 890 |
| *oxdC-rsiO* | 1274 | 644 | 11113 | 1152 | 339 | 577 |
| *cydABCD* | 73 | 39 | 12484 | 22 | 138 | 18 |
| *ldh-lctP* | 697 | 297 | 58791 | 180 | 1484 | 474 |

[1] The numbers indicate the intensities as determined by the transcriptome analysis. For operons, data for the first gene are shown. The data are sorted according to the ratio of expression in medium containing ammonium and 0.1 mM potassium for the wild type 168 vs. the Δ*dac* strain GP2222.

**S2 Table.**

**Transcriptomic data of genes involved in c-di-AMP, potassium, and glutamate homeostasis.** [1]

| Gene name | Wild type | | | | Δ*dac* | |
|---|---|---|---|---|---|---|
| | Glutamate | | Ammonium | | Ammonium | |
| | 0.1 mM KCl | 5 mM KCl | 0.1 mM KCl | 5 mM KCl | 0.1 mM KCl | 5 mM KCl |
| c-di-AMP homeostasis | | | | | | |
| *cdaA* | 10727 | 3014 | 6228 | 5878 | 18 | 20 |
| *disA* | 7600 | 2513 | 2492 | 3003 | 15 | 17 |
| *cdaS* | 26 | 621 | 30 | 19 | 17 | 19 |
| *pgpH* | 6932 | 2678 | 2860 | 3549 | 6659 | 4696 |
| *gdpP* | 4593 | 2199 | 6286 | 3623 | 6064 | 4994 |
| Potassium homeostasis | | | | | | |
| *kimA* | 98837 | 884 | 49463 | 2988 | 60648 | 7702 |
| *ktrA* | 6534 | 227 | 2462 | 645 | 8091 | 1662 |
| *ktrB* | 6425 | 301 | 1329 | 732 | 6573 | 1600 |
| *ktrC* | 3838 | 3294 | 2117 | 4643 | 5309 | 9220 |
| *ktrD* | 2729 | 1823 | 1248 | 1530 | 3246 | 2209 |
| Glutamate metabolism | | | | | | |
| *gltA* | 1529 | 2373 | 9269 | 59982 | 32726 | 62696 |
| *gltB* | 1917 | 2107 | 14529 | 69399 | 41312 | 71655 |
| *gltT* | 13499 | 6714 | 5335 | 12208 | 10594 | 12087 |
| *gltP* | 69 | 698 | 81 | 247 | 90 | 381 |
| *aimA* | 9882 | 1846 | 8135 | 10972 | 18750 | 17509 |

[1] The numbers indicate the intensities as determined by the transcriptome analysis.

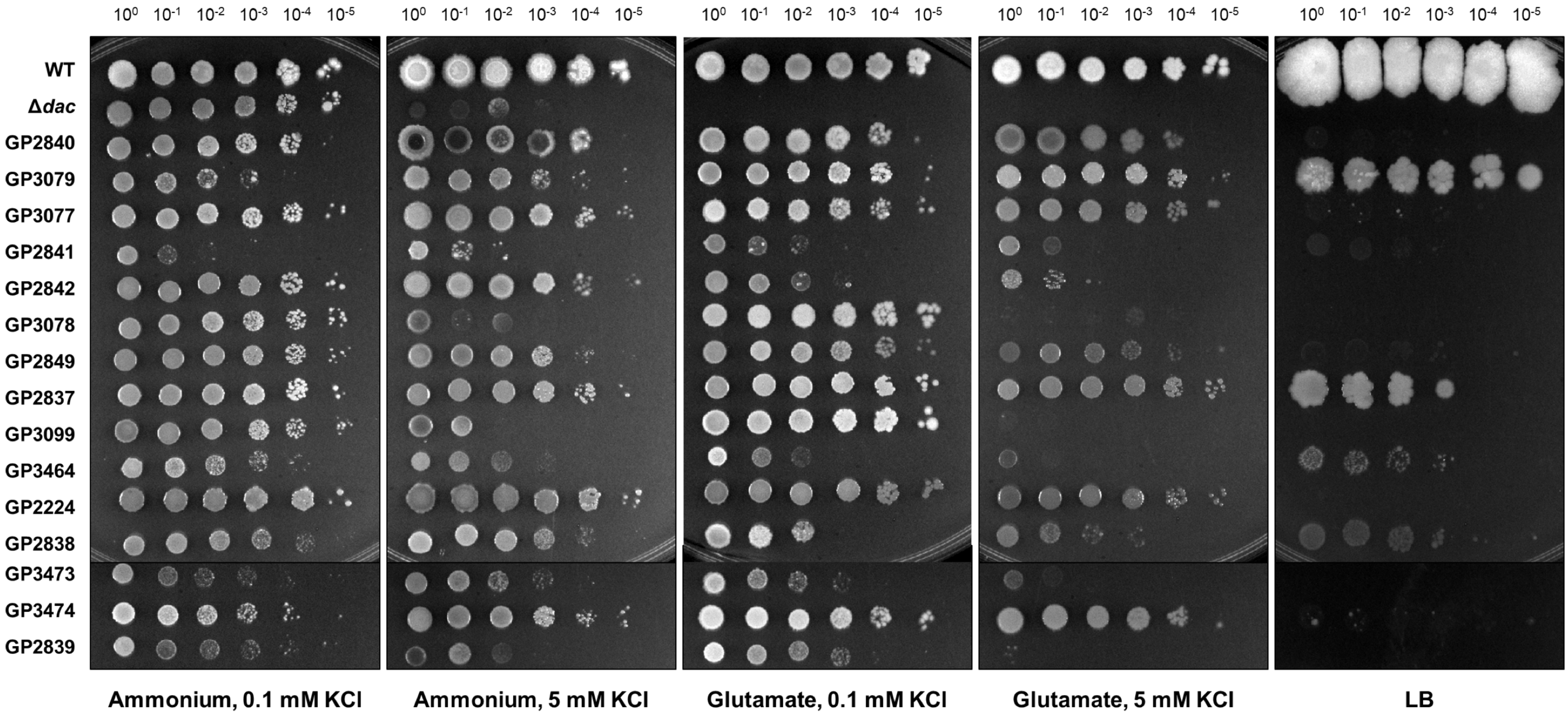


**S1 Fig. Acquisition of beneficial mutations allows growth of a c-di-AMP deficient strain in the presence of glutamate.**

Growth assay of *B. subtilis* wild type, GP2222 (Δ*dac*), and the isolated glutamate suppressor mutants. *B. subtilis* strains were cultivated in MSSM minimal medium with 0.1 mM KCl and ammonium. The cells were harvested, washed, and the $OD_{600}$ was adjusted to 1.0. Serial dilutions were dropped onto MSSM minimal plates with the indicated potassium concentration and ammonium or glutamate, or on LB plates.

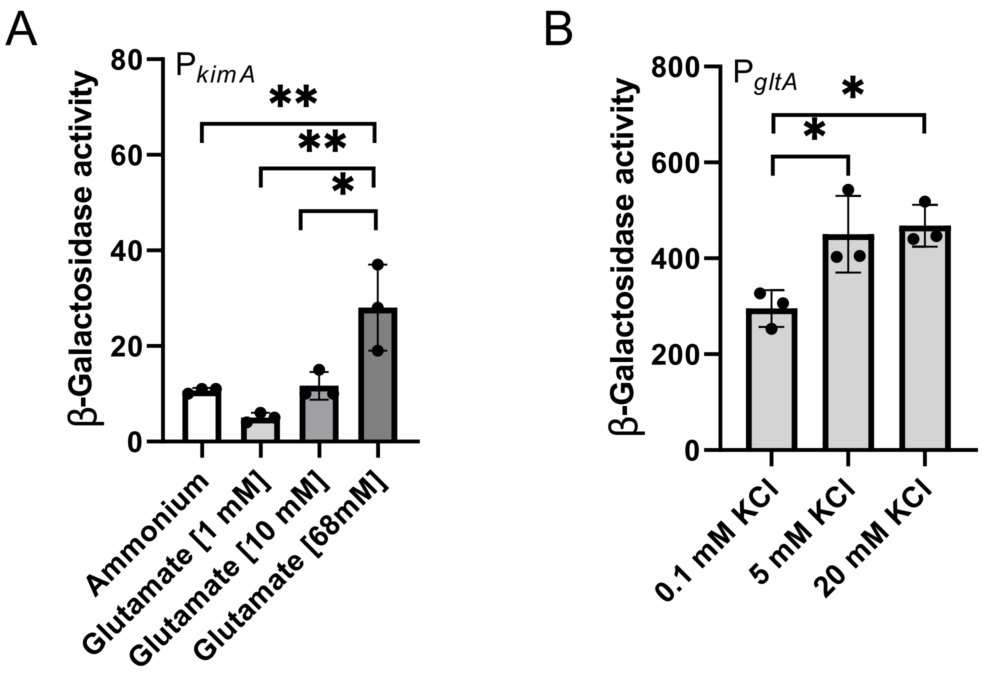


**S2 Fig. Potassium and glutamate influenced promoters.**

The expression of the high-affinity potassium transporter *kimA* (A) and the glutamate synthase *gltA* (B) was assessed by fusion of the promoter to the reporter gene *lacZ*. The cells harboring the promoter-fusion were cultivated with ammonium or glutamate at 0.1 mM KCl (*kimA*) or ammonium and different potassium concentrations (0.1, 5, 20 mM; *gltA*). Promoter activity was analyzed by quantification of β-galactosidase activity. Statistical analysis was performed using a one-way ANOVA, followed by Tukey's multiple comparisons test (**** $P < 0.0001$).

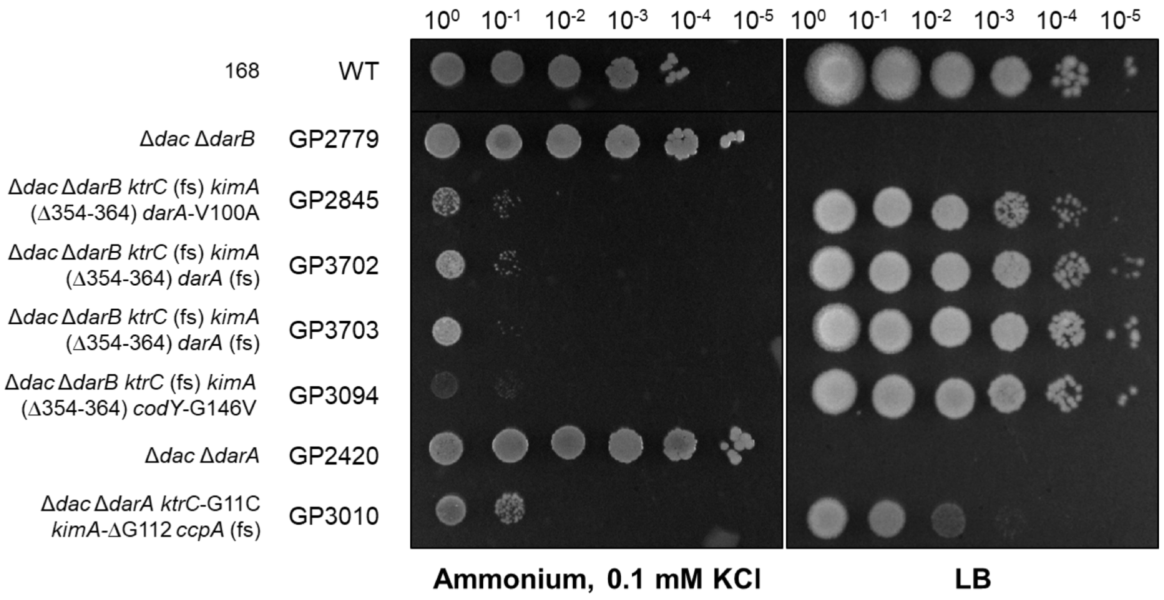


**S3 Fig. Two-step adaptation to complex medium.**

Growth assay of *B. subtilis* wild type, GP2779 (Δ*dac* Δ*darB*), GP2420 (Δ*dac* Δ*darA*) and the isolated glutamate suppressor mutants. B. subtilis strains were precultivated in LB medium. The cells were harvested, washed, and the $OD_{600}$ was adjusted to 1.0. Serial dilutions were dropped onto MSSM minimal plates with 0.1 mM KCl and ammonium or on LB plates.

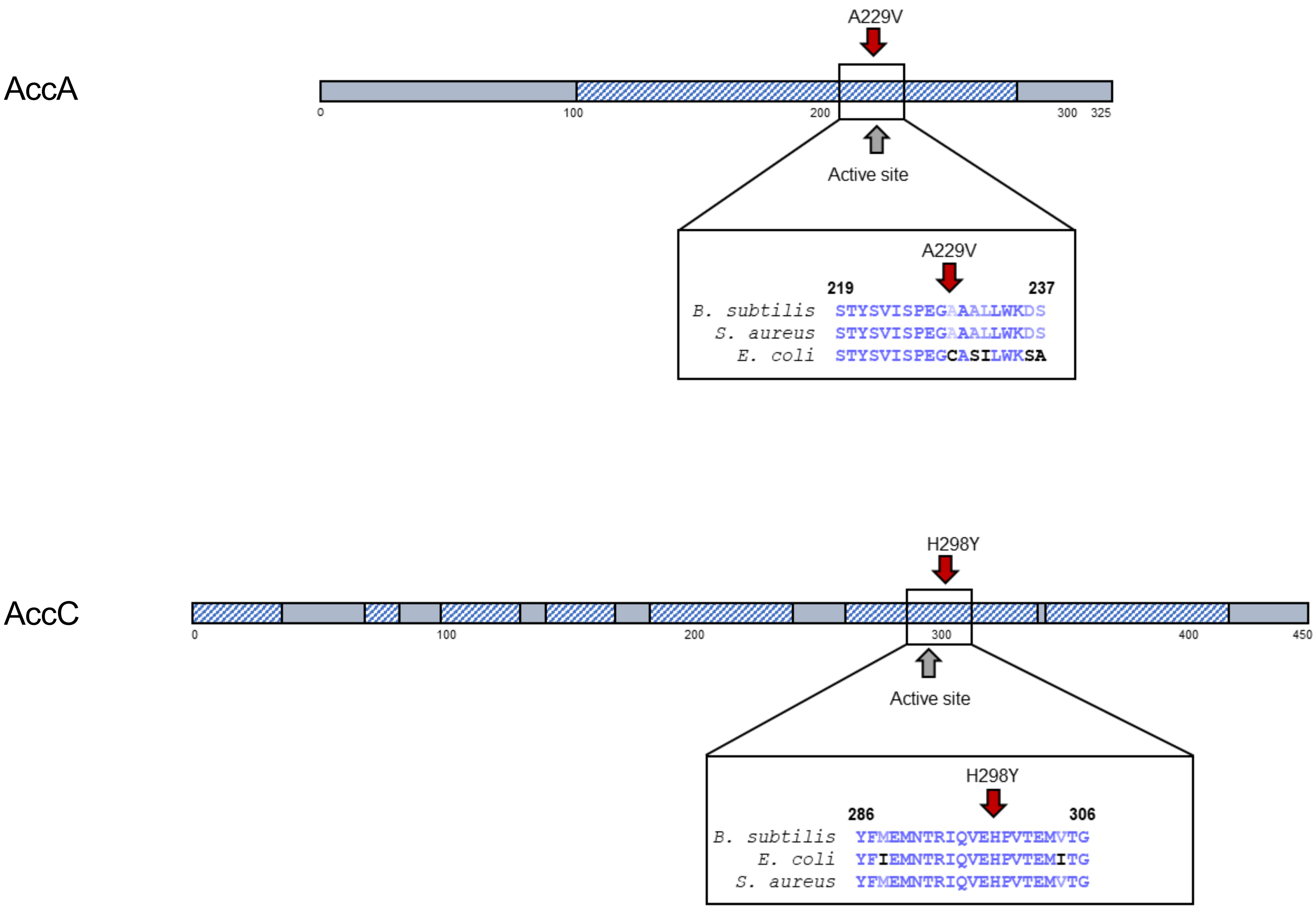


**S4 Fig. Mutations in the acetyl-CoA carboxylase subunits.**

Mutations obtained in the suppressor screen with glutamate (Table 1) and LB (Table 4). All mutations were single amino acid substitutions from independently isolated clones. Both mutations are located in highly conserved regions of AccA and AccC (hatched boxes) close to the active center.

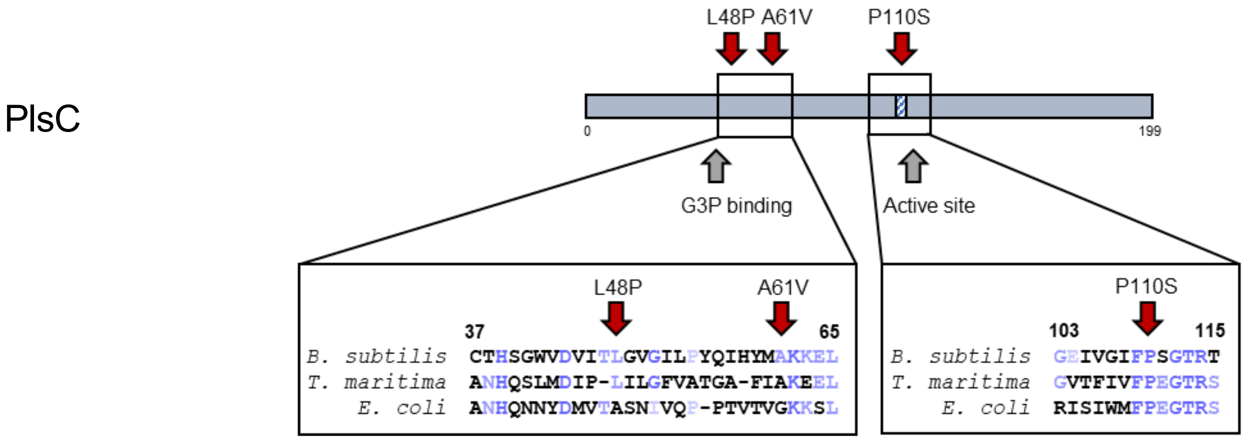


**S5 Fig. Mutations in the acyl-ACP:1-acylglycerolphosphate acyltransferase.**

Mutations in the *plsC* gene were obtained in the suppressor screen with glutamate (Table 1). All mutations were single amino acid substitutions from independently isolated clones. All mutations are located in functionally important regions of the enzyme.

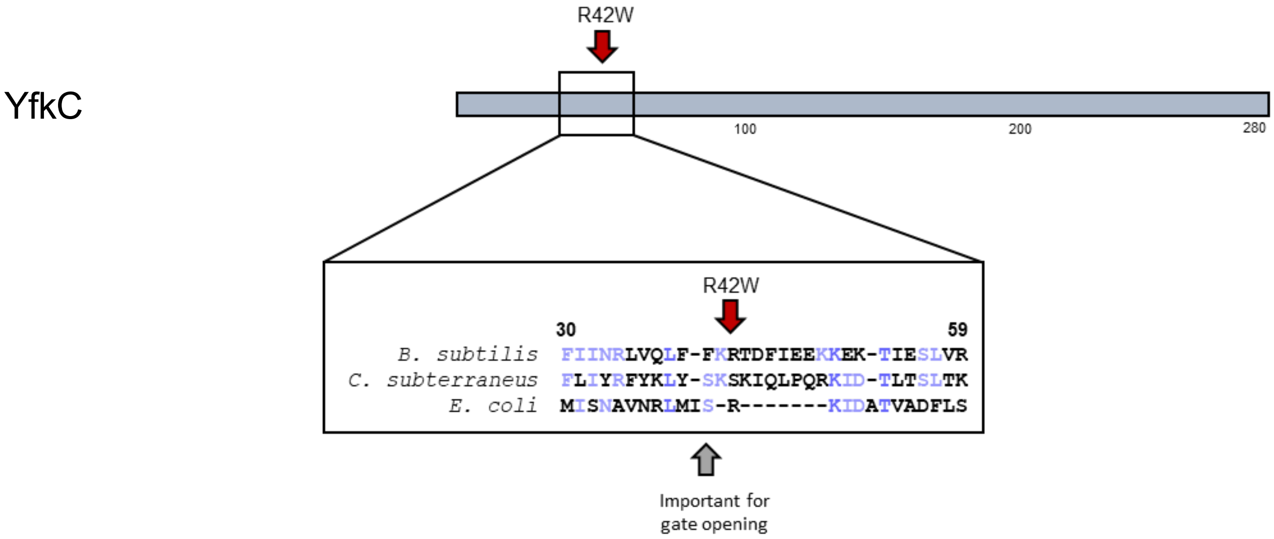


**S6 Fig. Mutations in the mechanosensitive channel YfkC.**

Mutations in the *yfkC* gene were obtained in the suppressor screen with glutamate (Table 1). The amino acid substitution Arg42 to Trp was observed in three independently isolated clones. In *E. coli* localization of S58 was observed to be highly dependent on the closing state of the channel (Miller *et al.*, 2003), thus, it appears likely that amino acid residue 42 in *B. subtilis* is located close to the gate.

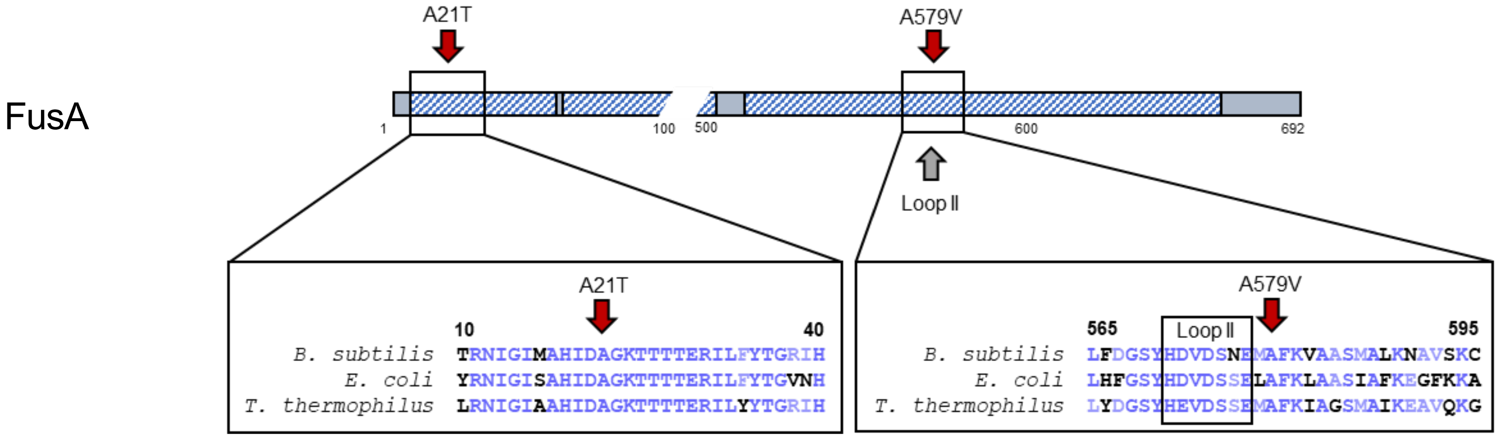


**S7 Fig. Mutations in the elongation factor G (FusA).**

Mutations in the *fusA* gene were obtained in the suppressor screen of GP3054 (Δ*dac* Δ*aimA*) with glutamate (Table 3). The amino acid substitution Ala21 to Thr was observed in one suppressor. Ala21 is adjacent to Asp20, an amino acid residue that binds the Mg ion and is therefore crucial for GTPase activity of the protein (Maracci *et al.*, 2014). In the two other suppressors Ala579 was mutated to Val. This amino acid residue is located in the conserved loop II that is required for proper translocation (Peng *et al.*, 2019).

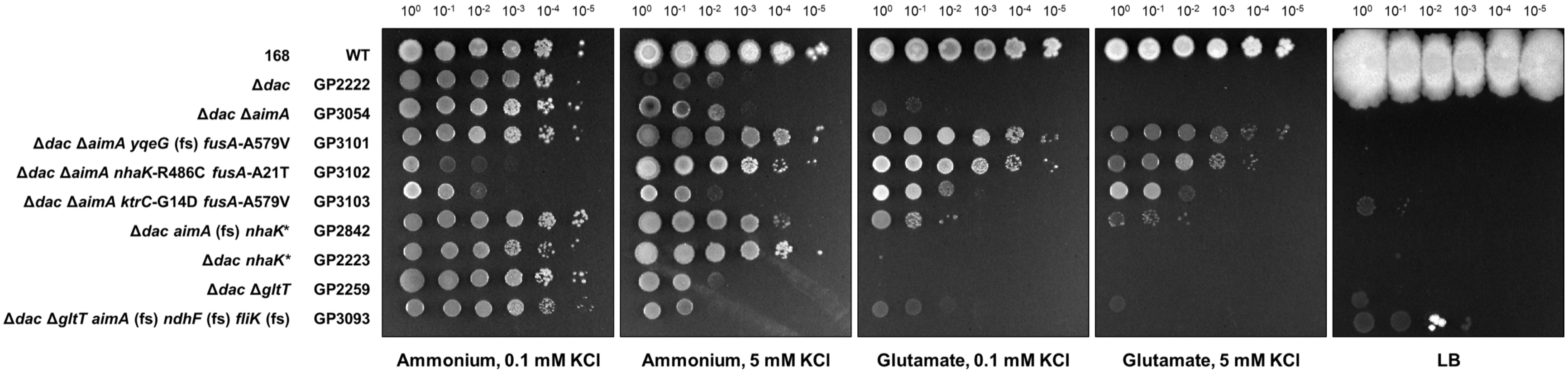


**S8 Fig. Effect of the deletion of the glutamate transporters in a c-di-AMP deficient strain.**

Growth assay of *B. subtilis* wild type, GP2222 (Δ*dac*), GP3054 (Δ*dac* Δ*aimA*), GP2248 (Δ*dac* Δ*gltT*) and the isolated glutamate suppressor mutants. *B. subtilis* strains were cultivated in MSSM minimal medium with 0.1 mM KCl and ammonium. The cells were harvested, washed, and the $OD_{600}$ was adjusted to 1.0. Serial dilutions were dropped onto MSSM minimal plates with the indicated potassium concentration and ammonium or glutamate, or on LB plates.
